# Combining radiomics and mathematical modeling to elucidate mechanisms of resistance to immune checkpoint blockade in non-small cell lung cancer

**DOI:** 10.1101/190561

**Authors:** Daryoush Saeed-Vafa, Rafael Bravo, Jamie A. Dean, Asmaa El-Kenawi, Nathaniel Mon Père, Maximilian Strobl, Charlie Daniels, Olya Stringfield, Mehdi Damaghi, Ilke Tunali, Liam V. Brown, Lee Curtin, Daniel Nichol, Hailee Peck, Robert J. Gillies, Jill A. Gallaher

## Abstract

Immune therapies have shown promise in a number of cancers, and clinical trials using the anti-PD-L1/PD-1 checkpoint inhibitor in lung cancer have been successful for a number of patients. However, some patients either do not respond to the treatment or have cancer recurrence after an initial response. It is not clear which patients might fall into these categories or what mechanisms are responsible for treatment failure. To explore the different underlying biological mechanisms of resistance, we created a spatially explicit mathematical model with a modular framework. This construction enables different potential mechanisms to be turned on and off in order to adjust specific tumor and tissue interactions to match a specific patient's disease. In parallel, we developed a software suite to identify significant computed tomography (CT) imaging features correlated with outcome using data from an anti-PDL-1 checkpoint inhibitor clinical trial for lung cancer and a tool that extracts these features from both patient CT images and “virtual CT” images created from the cellular density profile of the model. The combination of our two toolkits provides a framework that feeds patient data through an iterative pipeline to identify predictive imaging features associated with outcome, whilst at the same time proposing hypotheses about the underlying resistance mechanisms.

## I. INTRODUCTION

Lung cancer is the leading cause of cancer death in the U.S. for both women and men [1]. While cancer survival overall has improved, reduction in lung cancer mortality has been modest [1]. This may be attributed to late diagnosis, early metastasis, and development of resistance to conventional and targeted therapies. Accordingly, efforts had been made to apply different therapeutic modalities to the treatment of lung cancer. Immunotherapies, in particular, have had a profound and durable response in a variety of other metastatic cancers [2]. In 2016, checkpoint inhibitor immunotherapies were approved in the US as a first line therapy for some patients with non-small-cell lung cancer following the success of clinical trials using Nivolumab and Pembrolizumab [3]. Unfortunately, as has been seen with melanoma, even long term responders to these treatments may eventually acquire resistance [4], [5].

Checkpoint inhibitors are monoclonal antibodies that enhance the immune response to cancer by blocking inhibitory signals that restrict T-cell cytotoxicity. T-cells are an important part of the adaptive immune system that helps fight “foreign” cells, such as cancer cells. However, cancer cells can evade the immune attack by downregulating T-cell activity. One specific pathway that checkpoint inhibitors can target is the PD-1/PD-L1 pathway, which acts as an “on/off” switch for immune activity. When PD-1, a checkpoint protein on tumor-reactive T-cells, binds to PD-L1, a protein usually expressed on macrophages and some cancer cells, T-cell reactivity is turned off. Therefore, checkpoint inhibitors (anti-PD-1 or anti-PD-L1 therapeutic antibodies) can boost the immune response by disrupting the interaction between these cell surface proteins [6]. Expression of PD-L1 in tumor cells (or T-cells) or PD-1 in tumor infiltrating T-cells are associated with a larger likelihood of response to checkpoint inhibitors, although the predictive power of these biomarkers is not compelling [7], [8]. Some clinical trials don’t even take into account a significant threshold for PD-L1 expression [9]. Furthermore, the spatial context and dynamics of PD-L1 expression in both the tumor and inflammatory microenvironment is often ignored [9], [10].

In addition to intratumor heterogeneity in PD-L1 status, other microenvironmental factors may cause treatment failure. Heterogeneous and inefficient vasculature, contributes to regions of hypoxia and acidosis [11], [12]. In turn, acidity and hypoxia can block T-cell activation and induce severe anergy [13]. The study also found that neutralizing acidity in combination with checkpoint inhibitors elicited a synergistic anti-tumor response.

Radiomics attempts to quantify features from radiological images and use them as predictive or prognostic biomarkers of treatment response. These features are used as covariates in statistical or machine learning models to predict individual patient outcomes, and have been utilized to predict treatment response from pre-treatment images in multiple contexts [14], [15]. Statistical radiomics models enable correlations between imaging features and treatment response to be inferred, however, they cannot provide mechanistic explanations for the causes of treatment success or failure. This is beginning to change, however, as a recent report has linked radiomic signatures in NSCLC to upregulation of inflammatory gene sets in a bi-clustering approach [16]. However, these associations are only now emerging and can benefit from biologically-informed mathematical modeling, which allows causal mechanisms of treatment response to be explored.

In a novel combination of mathematical modeling with radiomics, we developed a spatial model of tumor cell and T-cell proliferation focused on response to immunotherapy. In parallel, we trained a radiomics model using computed tomography (CT) imaging data from lung cancer immunotherapy patients. We used these tools to gain insight into the mechanisms of treatment resistance. To link the cell-resolution mathematical model to the millimeter resolution radiological images, we developed a method to generate “virtual CT” images from the mathematical model and compare the features derived from the virtual CT images to the features identified in the patient CT images. The tools that we have developed enable an exploration of multiple mechanisms of immunotherapy resistance that can generate new hypotheses to guide future experiments.

## II. RADIOMIC FEATURES ASSOCIATED WITH TREATMENT RESPONSE

CT image data were available for a cohort of 51 metastatic lung adenocarcinoma patients treated as part of three different checkpoint blockade clinical trials conducted at Moffitt Cancer Center. The CT data consists of a 3-dimensional map of Hounsfield units (HUs), a measure of radiodensity. Radiomic features such as intensity, shape and texture were computed using methods described previously [17], [18]. Sparse partial least-squares-discriminant analysis was performed to determine the radiomic features with strongest association with treatment response. Response was defined as good (complete or partial response) versus poor (stable or progressive disease) according to the RECIST version 1.1 criteria. Figure 1 shows the loading weights of the first partial least squares component for the five features with strongest association with treatment response. We found that the skewness (a measure of the asymmetry of the distribution) of the HU histogram was most strongly associated with outcome, followed by the root mean square of the HU distribution, relative volume of air in the segmented tumor, then mean and median of the HU distribution.

**Figure 1.**
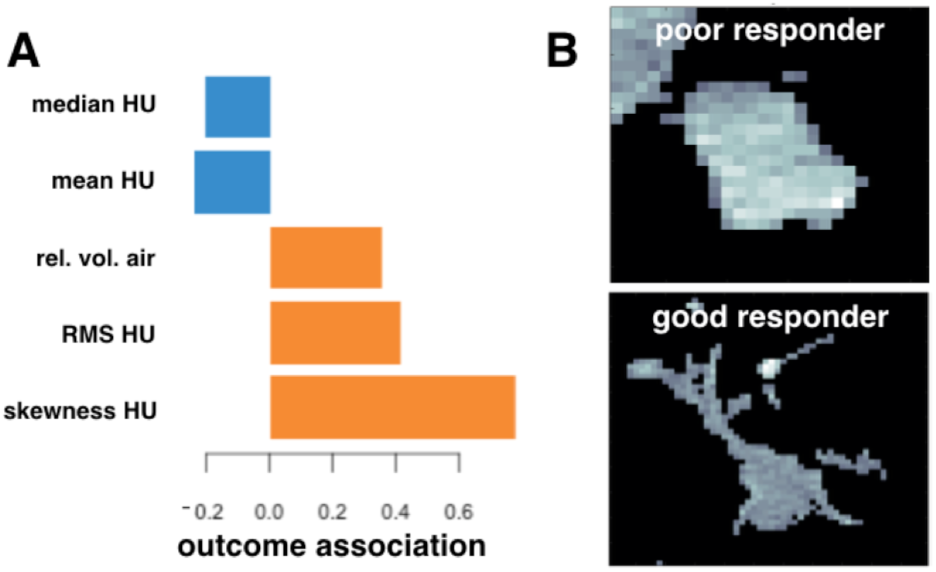
Results of the radiomics analysis. A) Sparse partial least squares discriminant analysis loadings of the 5 variables with strongest association with treatment response. Orange bars indicate that an increase in the feature value is associated with response to treatment. Blue bars represent that an increase in the feature value is associated with non-response to treatment. B) Example of a CT image of a lung tumor with poor response (stable or progressive disease) and good response (partial or complete response) to immunotherapy.

We also developed a computational toolbox to extract radiomic features concerning the shape and texture of the tumor from a subset of images from the cohort. This tool is equally capable of performing the same analysis on results from our mathematical model for comparison. Response to immunotherapy was found to correlate negatively with the tumor convexity and positively with the edge-to-core size ratio. The images in Fig. 1B show one tumor that responds poorly and one that responds well. While higher convexity (more round) has been shown in lung tumors to correlate with better survival [14], it is an interesting finding here that the more concave, invasive tumors respond better to immunotherapy. It is not clear why this occurs, but a possible explanation is that the more aggressive tumors that respond worse to conventional therapies are more susceptible to immune modulation. In a recent study patients who had faster growing tumors before initiation of immunotherapy had better responses [19]. To get a better perspective of the mechanisms involved, we developed a mathematical model to describe how checkpoint inhibition can affect tumor cells and their microenvironment.

## III. MODULAR MODEL OF RESISTANCE MECHANISMS

Our model considers spatial interactions of different types of cells through a number of biological processes projected to be relevant. Most of these processes were implemented as separate modules, which can be turned on and off, allowing for a mutable model with the ability to test the impact of adding or removing different elements. The intention is to elucidate how mechanisms, or combinations of mechanisms, affect patient outcomes.

The system was modeled as a spatial simulation of the tumor microenvironment with a stochastic partial differential equation based structure. We considered four main populations that compete for resources within a two-dimensional plane: *normal cells*, which constitute the untransformed epithelial tissue and stromal cells; *PD-L1 cells*, which are those cancerous cells that may successfully bind to the PD-1 receptors of the T-cells; *non-PD-L1 cells*, comprising the malignant cells that lack sufficient immune checkpoints on the cell surface to evade attack; and *glycolytic* cells, which produce diffusible immune-inhibiting factors as a byproduct of glycolytic metabolism. All cells have a turnover, but cancer cells proliferate faster than normal cells, with a small cost to proliferation associated with producing PD-L1 and using glycolysis.

In order to survive and proliferate, all cells require nutrients which are introduced into the system via randomly distributed blood vessel entry points and are further dispersed via diffusion. The local oxygen concentration modulates the proliferation rates of cells. The immune system’s T-cells enter through the blood vessels and interact with the cancer cells, destroying non-PD-L1 cells, but are simultaneously suppressed by their interaction with PD-L1 cells. Finally, the drug is introduced via the blood vessels and reverses the T-cells’ suppression by PD-L1 cells upon contact. The full interaction network of these mechanisms is shown in Fig. 2. Using the modular model setup, we can investigate each mechanism separately and combined to explore how treatment response could manifest both spatially and temporally.

**Figure 2.**
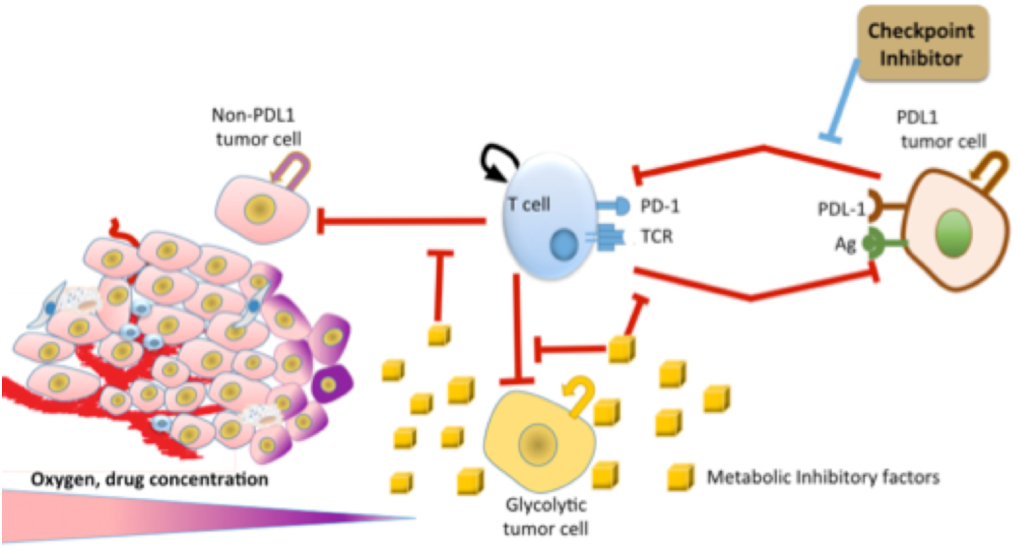
We consider both tumor and microenvironmental factors in the model. Tumor cells are either PD-L1 positive, PDL1 negative, or glycolytic, which affects response to T-cell kill. Growth and immune interactions are also modulated by spatially-dependent concentrations of oxygen, drug, andmetabolic inhibitory factors.

## IV. PD-L1 STATUS AFFECTS TUMOR SHAPE AND RESPONSE

To demonstrate how the interaction between the tumor and its microenvironment manifests at the spatial imaging scale, we investigated a simple example within this framework consisting of a tumor with only PD-L1 and non-PD-L1 cancer cells. We ran the model with two different compositions of cell types and otherwise identical initial conditions. One tumor was composed of a greater proportion of non-PDL1 tumor cells to PDL1 tumor cells, and in the second the proportions were reversed.

Figure 3 shows the population dynamics of the two tumors and the spatial distribution of cell types prior to treatment. The density profile shown is representative of a baseline clinical CT scan, which are commonly used to inform treatment planning. Regardless of the ratio of cell types, the dominant population outgrows and suppresses the minority population. The non-PD-L1 tumor (Fig. 3A) shows a more spiculated morphology, which can be explained by the fact that the non-PD-L1 cells are effectively attacked by the immune system. To proliferate and survive, the tumor must grow around blood vessels, which act as a delivery system for the immune cells. The PD-L1 cells, however, are able to evade immune attack; this facilitates their coexistence with the vasculature and allows them to form a more rounded mass (Fig. 3B).

**Figure 3.**
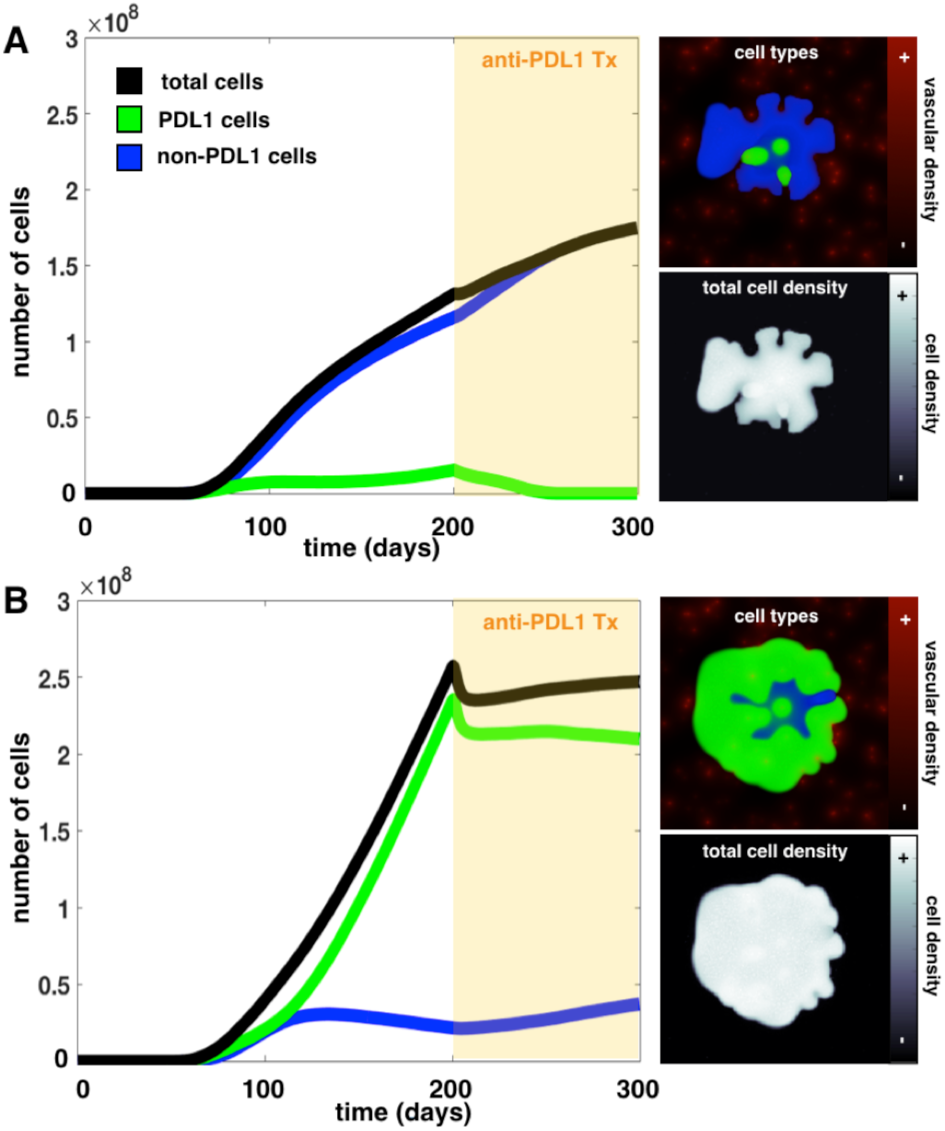
Comparing growth and treatment of a tumor that contains mostly non-PD-L1 cells (A) with a tumor that contains mostly PDL1 cells (B). Population dynamics plots show how each tumor composition grows and responds to anti-PD-L1 treatment. the spatial layout of cell types and total cell density is shown to the right of each plot.

When immunotherapy was introduced, the non-PD-L1 tumor effectively continued its growth trajectory. The death of the few PD-L1 cells was compensated by a smallincrease in the growth rate of non-PD-L1 cells. In contrast, the PDL1 tumor growth was significantly slowed as a result of treatment. With a larger number of targeted cells, there was a significant decrease in the PD-L1 population, which caused a decline in the total population of cells. After a period of response however, there was an increase in the total population; this was due to regrowth of the non-PD-L1 cell population after spatial competition with the PD-L1 cells was removed.

These results demonstrate that if tumor heterogeneity is based solely on PD-L1 status, deviation from isotropic invasion is a result of immune evasion by the non-PD-L1 cells, which lack the PD-L1 suppression mechanism. However, using shape from this model as a predictor for response is in exact opposition to the data. It is clear that more factors are involved than just cell type and that this mechanism may not be the most dominant factor in determining tumor response. We can, however, further characterize the tumor profile using the cell density matrix as a “virtual image” and extract radiomics features. We quantified several features from the virtual images that overlapped with the features with the highest outcome associations from the exploratory radiomics analysis and found that some features agreed with the data (RMS HU and skewness HU), and others did not (convexity and mean HU). It is possible that these features match well with the model because they correspond to intratumor heterogeneity characteristics, which we investigated here by using different ratios of cell types. At the same time, other mechanisms not considered here may be stronger drivers of shape and overall density.

## V. DISCUSSION

Quantitative radiological features extracted from routine patient imaging have been shown to correlate with patient outcome [14], [15]. Often clinical radiological assessment is based on the size and number of tumors alone, while the tumor heterogeneity, shape, and dynamics are observed but not quantified. In our analysis, we found several radiomics features correlated with outcome using a cohort of patients with heterogeneous response to immunotherapy, and found a single mechanism that underlies changes shape and texture characteristics of tumors. The relationship between tumor features and molecular or phenotypic characteristics of the tumor and the microenvironment are complex. However, we propose that relating feature to function can be possible through mathematical modeling.

The computational structure of this mathematical model was designed to easily turn on and off individual components to enable investigation of how each might contribute to overall tumor growth and treatment response. Mathematical models of biological phenomena can often be overly complex, leading to overfitting to available data and inability to make accurate predictions. The basis of modeling is to simplify a system in order to understand it. However, when dealing with large heterogeneous populations of cells that interact with each other and their environment, overly simple models often neglect relevant biological processes. Our computational framework was created to be both reductionist and integrated, in order to try to understand both the singular components and the interconnected whole.

The main portion of this work was developed over an intense 4-day workshop and is just the start of a broader framework to use mathematical modeling to bridge cellular and tissue level mechanisms to large cohorts of patient data. Statistical analysis of clinical data, histological features, and imaging data are regularly used to define prognostic and diagnostic criteria, but the focus is usually on accurately determining patient outcome rather than the mechanisms leading to those outcomes.

In this work we proposed a novel framework for bridging multiscale data by combining two approaches that have previously been applied separately: statistical radiomics modeling and mechanism-driven mathematical models. Statistical radiomics modeling enables identification of imaging features that correlate with patient response to currently employed treatment strategies and the prediction of individual patient outcomes. Mathematical modeling, on the other hand, through the quantitative description of biologically-relevant mechanisms, may be used to test many different treatment strategies computationally. Promising treatment strategies informed by the mathematical models can then be validated using a range of biological experiments. Our framework uses multiscale data and combines both approaches. The use of all available data and multi-disciplinary analysis feeds-forward a refined understanding of underlying biological interactions and a narrowed set of radiomic features that are correlated to mechanisms of interest for treatment success. Using a larger cohort of patient data would allow for responders and non-responders to be better defined on all scales so that the range of possible responses for a specific tumor can be reduced. Testing different treatments on the reduced “model space” can then be assessed for the best and most probable outcome.

**Figure 4.**
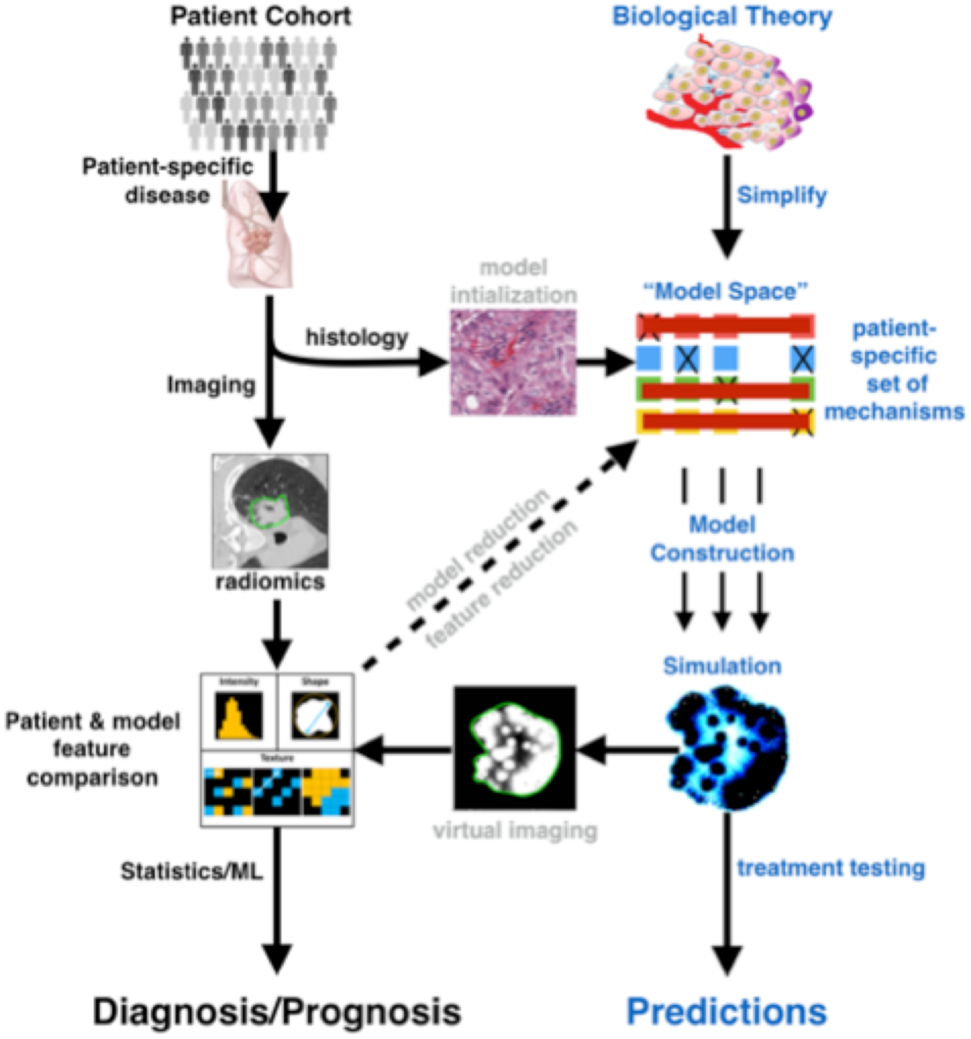
Data-driven, statistical-based radiomics approaches and mechanism-based mathematical models are often used for separate purposes. We propose a larger framework that bridges patient data to biological mechanisms by using tools from each approach to iteratively inform the other.

## VI. ACKNOWLEDGMENT

We would like to thank the IMO Chair, Dr. Alexander Anderson, for organizing the 6th Annual Moffitt IMO workshop: Resistance, where this project was conceived. We are also extremely grateful to the Moffitt Cancer Center and the Moffitt PSOC for supporting this workshop through the NCI U54CA193489 grant. Thank you also to Matthew Schabath for curation of the trial data and Anthony Magliocco and Mark Robertson-Tessi for their insightful discussions.

## Notes

Research supported by H. Lee Moffitt Cancer Center & Research Institute.

